# Continuous site-directed mutagenesis and selection in *Escherichia coli*

**DOI:** 10.64898/2026.07.18.739322

**Authors:** Fabian B. H. Rehm, Martin Spinck, Jason W. Chin

**Affiliations:** Medical Research Council Laboratory of Molecular Biology, Francis Crick Avenue, Cambridge, England, UK; Generative Biology Institute, Ellison Institute of Technology, Heatley Rd, Oxford, England, UK

## Abstract

The programmed diversification of genes and other encoded genetic elements, through site directed- and site saturation-mutagenesis, underpins approaches to learning the relationship between DNA sequence and function, and forms a foundation for creating new function through directed evolution. However, current approaches for generating large genetic libraries commonly generate diversity *in vitro* and then transform the resulting library into cells; this multistep process is inefficient and this paradigm places limits on the scale of diversity that can be achieved and the size of diversified genetic elements that can be introduced into cells. Here we demonstrate a distinct paradigm for library generation through: 1) efficient transfer of genetic elements, as cargos in F plasmids, to recipient cells bearing libraries of retron editors, 2) efficient editing of cargo genes in recipient cells, and 3) continuous iteration of the conjugation-editing cycles with selection for recipients in sequential cycles using three selection markers in series. In this paradigm, the library diversity emerges multiplicatively through the iteration of conjugation-editing cycles. Using this paradigm, we generated substantial libraries that enabled the selection of new phenotypes, with library members containing up to six distinct edits and edits arising from several conjugation-editing cycles.

---

The programmed diversification of genes and other encoded genetic elements, through site directed mutagenesis, underpins approaches to learning the relationship between DNA sequence and function, and forms a foundation for creating new function through directed evolution^1^. The majority of site directed mutagenesis approaches operate *in vitro* and generate diversity by rounds of oligonucleotide directed mutagenesis^2^. In these approaches, libraries are commonly created by cycles of directed mutagenesis with the overall diversity resulting from the product of the diversity introduced by the pools of mutagenic oligonucleotides used in each cycle. For experiments in cells, the resulting DNA mutants or mutant libraries need to be transformed and this adds an additional step that can limit the transfer of large libraries generated by *in vitro* methods, and scalability for large scale studies^3^. These challenges are exacerbated further with larger genetic elements.

While a variety of *in vivo* editing methods facilitate site directed mutagenesis in cells^4–9^ the introduction of multiple edits is more challenging *in vivo* and commonly requires the iterative addition of editing systems, or their nucleic acid substrates, to cells containing the target for mutagenesis^10–12^, Because these approaches rely on delivering a functional editor to every cell and the editing provided by these systems is generally far from quantitative, it is not commonly possible to generate site directed libraries *in vivo* at the scale and depth that has been achieved by *in vitro* methods^13,14^. Moreover, repeated cycles of editing in the same cell often lead to off target mutations that alter the genetic background, and/or inactivate the editing system, further limiting the depth and scale of mutagenesis that may be accessed and therefore the sequence space that may be interrogated for function.

We have been interested in exploring a distinct paradigm for introducing combinatorial edits, or performing any other combination of operations, in a genetic element of interest (**Figure 1**) In this paradigm, a genetic element of interest is moved into a population of cells that each contain distinct members within a set of operators (eg: a set of editors that mutate a first target position within the genetic element to other options) and, these operators are allowed to operate on the genetic element to generate variation. The resulting population of varied genetic elements is then moved into a second population of cells, which contains a different set of operators (e.g.: a set of editors that mutate a second target position to other options), that are allowed to operate on the varied genetic elements generated in the first cells. The genetic element may be passed through many such cycles. The overall combination of operations is realised by distributing sets of operations between cycles and multiplying sets of operations by using the output of one cycle as the input for the next cycle. This approach is naturally suited to generating large libraries from small sets of operators, as the final library diversity is a function of the paths taken by the population of genetic elements through the libraries of operators in distinct cycles, rather than a function of the size of libraries that can be transformed into cells. This paradigm may enable the scalable site directed mutagenesis of genetic elements in cells. As this approach resets the cells genetic background in each cycle it inherently limits the effects of off target mutation on the operations performed.

**Figure 1:**
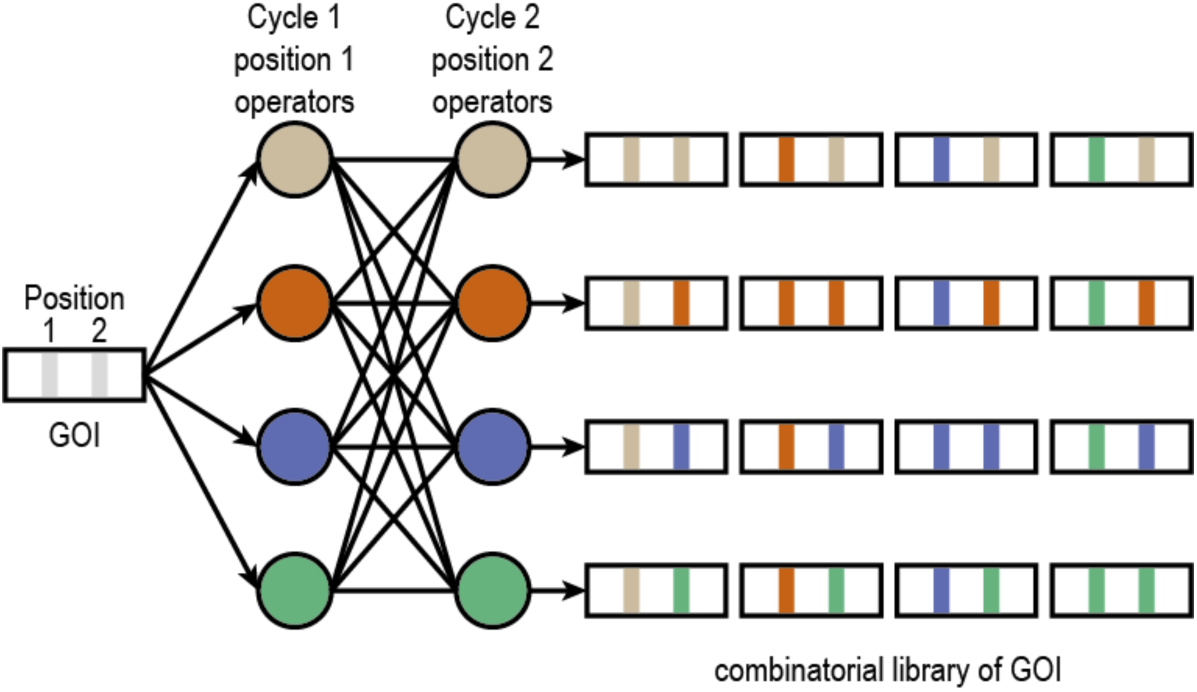
Generation of site-specific combinatorial libraries by driving a mobile cargo through a library of operator strains. A model highlighting the principle of a distributed approach to library generation. A genetic element (F-integrated cargo) with two positions of interest is shown. In the first cycle the genetic element is transferred to a library of operators (coloured circles) that each perform a different operation (e.g. editing) on position 1. In the second cycle the genetic element is transferred to a library of operators that each perform different operations on position 2. The path of each copy of a genetic element through the operators defines the modifications (coloured bars) accumulated in that copy. Distinct (random) paths of distinct copies of the genetic element lead to distinct combination of operations (edits) at positions 1 and 2. All combination of operations at position 1 and 2 may be achieved by the population of genes, sampling all possible paths through each library of operators.

Here we show that we can near-quantitatively transfer a genetic element of interest, as cargo in a self-transmissible F plasmid^15,16^, through a population of cells by conjugation. We establish retron editing systems with high efficiency^17^. We demonstrate that we can transfer a cargo genetic element through cells bearing a series of retron editors to efficiently generate a defined combination of site directed edits over a series of conjugation-editing cycles. We demonstrate that using three selection markers makes the conjugation-editing cycles scalable and repeatable over numerous cycles: the transconjugant cells for the *n*th conjugation-editing cycle act as the donor cells for the n+1 conjugation-editing cycle. Finally, we show that we can use our approach to generate diverse libraries of site-specific edits in a gene of interest, by transferring a cargo genetic element through several libraries of retron editors. This approach generates genetic elements that are diversified by several cycles and we exemplify the selection of a protein with a new phenotype from the resulting library.

## Results

### Efficient transfer of F-plasmid integrated cargo

To quantify the overall efficiency with which an F-integrated cargo is transferred from donor to recipient cells, we used a modified RK24 carrying an integrated 5.7 kb luciferase operon (*Photorhabdus luminescens* LuxABCDE); this F-plasmid confers luminescence to the cells carrying it (**Figure 2a**). We mixed an *E. coli* strain harbouring the RK24-lux F-plasmid or an inactive mutant lacking the origin of transfer (oriT) in a one-to-one ratio with a F-recipient (Cm^R^). Following conjugation, we determined the percentage of luminescent recipient (Cm-resistant) colonies. Overall, 65% ± 11.4% of recipient cells received the luciferase operon, confirming the highly efficient transfer of the F-integrated cargo from donor to recipient cells (**Figure 2b, Supplementary Figure 1**).

**Figure 2.**
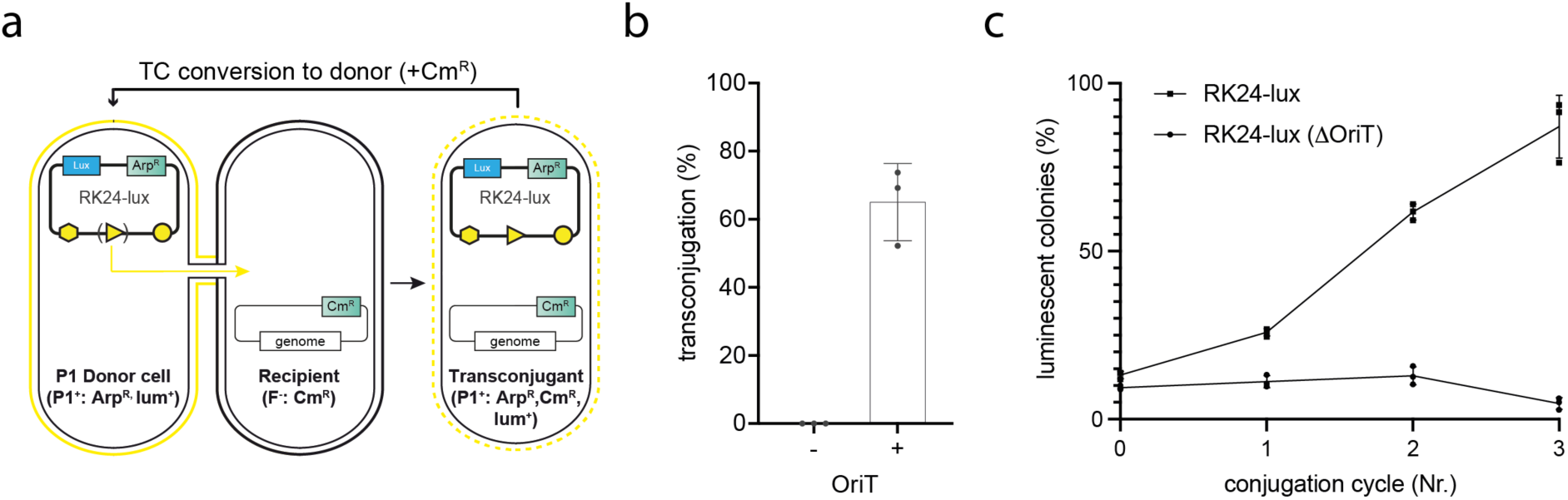
F plasmid with integrated cargo rapidly colonizes a recipient population. a) We used a luminescent version of the RK24 F-plasmid carrying the Lux operon of *Photorhabdus luminescens*. It also harbours the genes for conjugative transfer (yellow hexagon), an origin of transfer (OriT, yellow triangle) and machinery for replication and partitioning (yellow circle). The non-transferable version of RK24-lux is lacking the OriT. Conjugative transfer is initiated at the OriT, transferring the F-plasmid and integrated cargo into the recipient strain (Cm^R^). The recipient strain gains the full F-plasmid and the integrated markers and cargo. Shortly after conjugation the transconjugants convert into donor cells, allowing for repeated cycles of conjugation. b) Transconjugation efficiency of RK24-lux without and with OriT as a percentage of luminescent recipient cells following 1h of conjugation. The bars show the mean of three biological replicates with the error representing the standard deviation of the mean, individual datapoints are shown as black dots. c) Fraction of luminescent single colonies (as % total colonies) following three iterative cycles of conjugation of the self-transmissible (RK24-lux, black squares) or non-transmissible F-plasmid (RK24-lux ΔOriT, black circles). We started with mixed population containing 10% donor cells. A black line indicates the mean of three independent biological replicates and error bars show the standard deviation of the mean, individual datapoints are shown as black dots or squares.

### F-plasmid transfer through a bacterial population

Following conjugation of an F-plasmid from donor cells to recipient cells, the cells receiving the F-plasmid can act as donor cells. For F-plasmids carrying cargo, this should allow for the repeatable transfer of the F-integrated cargo through a population of recipient cells by conjugation. To demonstrate this transfer, we prepared a conjugation reaction containing a ten-fold excess of recipient to donor cells and iteratively cycled cells through conditions that favour conjugation (1 h on agar plates to allow sustained cell-cell contact) followed by 30 min of shaking in liquid medium (to break cell-cell contact and allow cell growth). Cells were then returned to agar plates to initiate a new cycle.

After three cycles the percentage of the population containing the F-plasmid carrying cargo increased from 13.2% ± 1.1% of the population to 87% ± 9.3% of the total population.

After each cycle, the percentage of the population containing the F-plasmid approximately doubled (1.9 ± 0.5-fold increase per cycle); thus, the fraction of transconjugant cells increases exponentially. As each cycle results in the conjugation of the F-plasmid carrying cargo from one donor to one recipient, we refer to these cycles as conjugation cycles. In control experiments, the percentage of the population containing a non-transmissible RK24-lux (ΔOriT) decreased from an initial population of 9.3% ± 0.6% to 4.7% ± 1.7% over three cycles (**Figure 2c, Supplementary Figure 2****and 3**).

**Figure 3.**
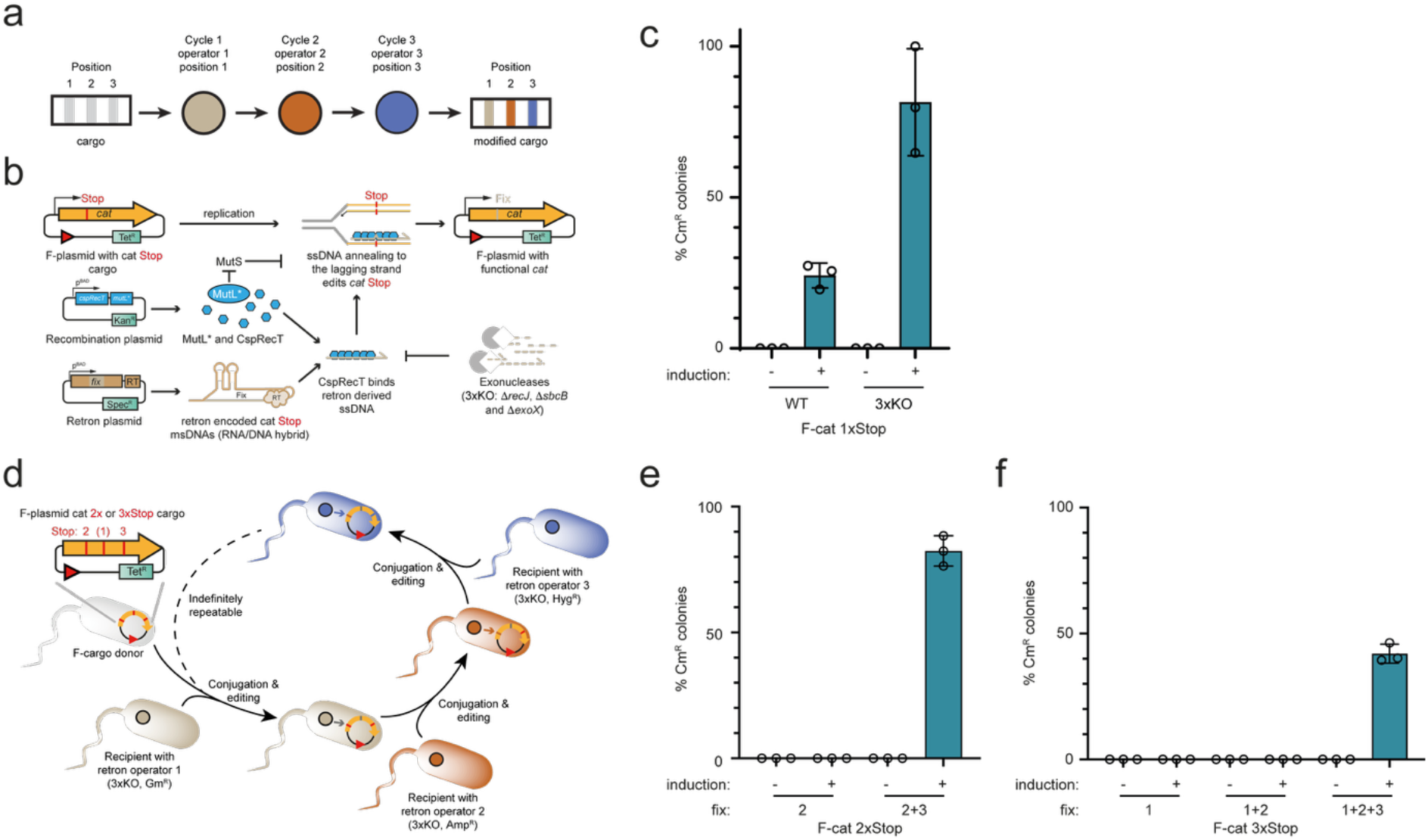
**Cycling F-integrated cargo through cells with efficient and distinct site-specific editors enables the accumulation of programmed edits**. a) Cycles of conjugation and editing allow for different operators to act on a mobile cargo, which accumulates designed site-specific edits defined by the operator used in each cycle. b) Schematic of F-integrated cargo editing via lagging strand annealing of a ssDNA as facilitated by CspRecT (blue hexagon). We reprogram the msDNA to reverse transcribe the target sequence via the associated retron reverse transcriptase (RT) generating intracellular ssDNA. The retron-derived ssDNA is bound by RecT which utilizes homology driven hybridization to the lagging strand intermediate during DNA replication to introduce site specific edits. Processes such as mismatch repair (MutS) or cellular exonucleases can decrease the editing efficiency by recognizing and repairing the mismatched lagging-strand intermediate or by degrading intracellular ssDNA, respectively. The dominant negative mutant MutL* (E38K) is known to repress mismatch repair and we generated an exonuclease knockout strain (3xKO), removing three non-essential exonucleases (RecJ, SbcB and ExoX). c) Fraction of chloramphenicol-resistant cells as a percentage of total cells obtained via CspRecT/MutL* based reversion of a single stop codon in a *cat* reporter utilizing a retron-derived ssDNA oligo, in DH10B (wt) or DH10B *ΔrecJ, ΔsbcB ΔexoX* (3xKO). Dark blue bars show the mean frequency of resistant cells with error bars showing the standard deviation of the mean, individual datapoints are shown as hollow circles. The experiment was performed in three biological replicates. d) Experimentally, we integrated a *cat* gene containing two or three stop codons on the pED208 F-plasmid (F-cat 2xStop or 3xStop) and transferred the cargo into a recipient strain containing the operator of choice (e.g. 3xKO strain with CspRecT/MutL* and retron encoded fix1, Kan^R^). The operator acts on the F-integrated cargo reverting the targeted stop codon and the F-plasmid converts the transconjugant cells (TC) into donor cells, these become the new donors for the next cycle with the next operator recipient strain. We rotate between three distinct recipient markers on the CspRecT/MutL* recombination plasmid, for more than three cycles this can be repeated indefinitely. e, f) Fraction of chloramphenicol resistance cells as a percentage of total cells containing the F-integrated cargo (*cat* 2xStop, e, or 3xStop, f) after each cycle of conjugation and editing by the respective operator strains in the presence and absence of arabinose induction of the retron-derived ssDNA and the recombination system. Dark blue bars show the mean frequency of resistant cells with error bars showing the standard deviation of the mean, individual datapoints are shown as hollow circles. The experiment was performed in three biological replicates.

Overall, these experiments demonstrated that an F-plasmid cargo in a small population of donor cells can be efficiently transferred into a large population of genetically identical recipient cells.

### Cumulative editing of an F-integrated cargo

Next, we demonstrated that we can perform distinct operations on the F-integrated cargo at each conjugation cycle by embedding distinct operators (which we defined as genetic elements that operate on the F-integrated cargo) within the recipient cells used in each cycle. We also demonstrated that after several conjugation cycles, the output is the product of the operations performed at each cycle (**Figure 3a**). We focussed on editing operations on the F integrated cargo to exemplify this paradigm.

Various strategies enable targeted gene editing in bacterial cells^4,18^. Although many editing strategies may be compatible with conjugation cycles, we focussed on identifying efficient editing methods. Efficiency of each editing step is important because the net editing efficiency across several editing cycles is product of the editing efficiencies for each cycle. We selected lagging strand annealing of retron-derived reverse transcribed ssDNA by the CspRecT/MutL* recombination system as it is well characterised and reasonably efficient (**Figure 3b**)^5^. To achieve high editing efficiency, we generated a DH10B *ΔrecJ ΔsbcB ΔexoX* knockout strain (DH10B 3Δ). Removing these three exonucleases from the genome improved single-step retron editing efficiency, over 3.5 h, from 24% to 81% (**Figure 3c**).

To demonstrate our approach, we generated F-plasmids (pED208 F-plasmid (cat 2xStop or cat 3xStop)) in which we integrated a chloramphenicol acetyl transferase gene (*cat*) containing two or three stop codons. We also designed retron-derived, arabinose-induced ssDNAs that target each stop codon within *cat2X stop* or *cat3X stop* and revert them to sense codons via a single base substitution.

We transferred the F-integrated *cat* 2xStop and *cat* 3xStop through editing strains with retrons targeting each stop codon in series (DH10B 3Δ, CspRecT/MutL* and inducible stop targeting msDNA), with the goal of reverting one stop codon in each cycle upon induction of the editing system.

Each conjugation-editing cycle consisted of a conjugative transfer step, followed by recovery and selection for transconjugants, and then a 4-6 h editing step before transfer into the next recipient carrying the next editor and a distinct resistance marker (**Figure 3d**). To enable repeated cycling, we used three different antibiotic resistance markers in recipients in alternate rounds. This marker strategy allowed for selection of transconjugants in each round with minimal carry-over from previous rounds^19^. Chloramphenicol resistance requires the conversion of all 2 or 3 stop codons to sense codons; thus, the cumulative editing efficiency over all steps was obtained by plating on chloramphenicol. The *cat* 2xStop cargo following two conjugation/editing cycles yielded 82±6% chloramphenicol-resistant cells, whereas the 3xStop cargo yielded 42±4% chloramphenicol-resistant colonies after three cycles (**Figure 3e,f**).

These experiments demonstrated that cycles of conjugation of an F-integrated cargo can be readily coupled to efficient *in vivo* editing of the cargo by a series of editors. This approach rapidly yields gene variants harbouring multiple site-specific edits.

### Continuous site-saturation mutagenesis via iterative conjugation-editing cycles

Next, we demonstrated that, by embedding a library of operators within the population of recipient cells used in each cycle (such that each recipient cell contains a distinct operator from within the library), we can perform a library of operations on the F-integrated cargo in each conjugation cycle. By using recipient cells bearing a different library of operators in each cycle, the output for each F-integrated cargo becomes a function of its path through the library of operators. We focussed on exemplifying this paradigm using libraries of retron editors that operate on positions in cargo genes to realise a site-directed mutagenesis library. In this approach, site-directed diversity in target genes is a function of the paths that the population of F-integrated cargos take through the population of cells bearing libraries of retron editors; thus, in this paradigm – unlike in most approaches to *in vivo* site directed library generation – diversity need not be bottlenecked by the transformation efficiency of cells.

We chose to exemplify this approach through the evolution of an aminoacyl-tRNA synthetase (aaRS) to accept non canonical amino acid (ncAA) substrates. We integrated the genes encoding pyrrolysine aaRS from *Methanococcus mazei* (*Mm*PylRS) and *Mm*PyltRNA_CUA_ onto the pED208 F-plasmid (**Figure 4a**). We targeted positions within the active site of PylRS using designed retron operator libraries that harboured degenerate NNK codons. We designed a total of five retron library sequences, four of these library sequences harboured one NNK while one library sequence harboured three NNKs (**Figure 4b**). As each of these libraries is small individually, there was no risk of library bottlenecking the accessible diversity due to transformation of the retron plasmids.

**Figure 4.**
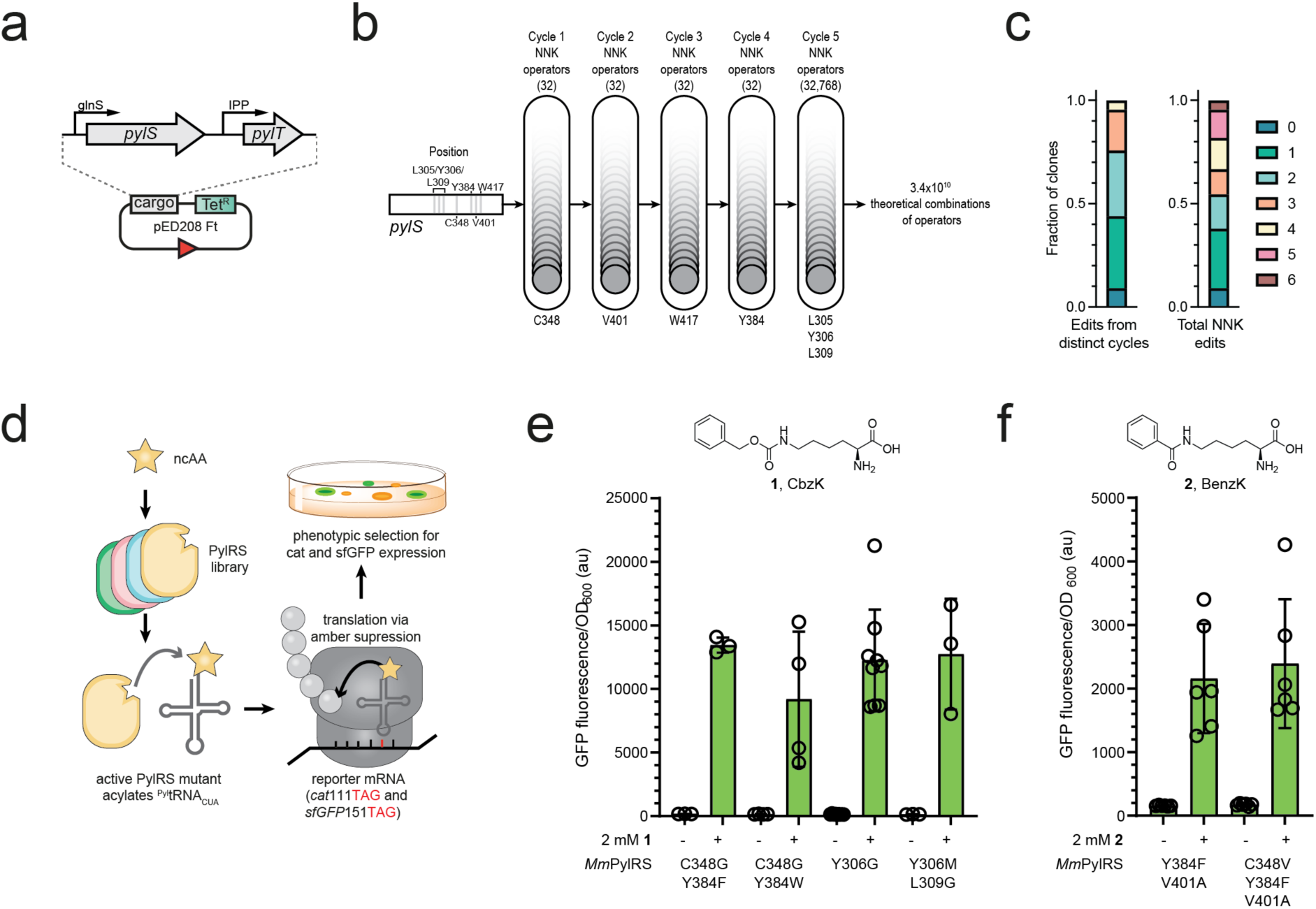
Cycling F-integrated cargo through cells with libraries of site-specific editors enables continuous and multiplicative cargo diversification and selection of new function. a) We integrated the *Mm*PylRS and *Mm*^Pyl^tRNA_CUA_ genes (PylS and PylT respectively) on the pED208 self-transmissible F-plasmid (via OriT mediated conjugative transfer, red triangle). b) We designed five retron operator libraries to site specifically install NNK degenerate codons at the indicated positions within *Mm*PylRS using the CspRecT/MutL* recombination system. The number of operators in each cycle is shown in brackets (an NNK library at a single position generates 32 operators). In each cycle the recipient selection markers are used in rotation (hgr^R^, kan^R^ and gen^R^). c) Fraction and number of edits arising from distinct editing cycles, and the total number of resultant NNK edits, observed in 66 randomly picked clones prior to any phenotypic selection. d) After the final cycle, the resulting library was transferred from the editing strain into a selection strain (carrying *cat*112TAG, *sfGFP*151TAG, reporter plasmid). Active PylRS mutants were selected for GFP fluorescence and chloramphenicol resistance in the presence of ncAAs (CbzK or BenzK). e,f) GFP expression in the presence or absence of the ncAA CbzK (1, f) or BenzK (2, g). Bar graphs show the mean GFP fluorescence, with the error bars indicating the standard deviation of the mean, individual datapoints are shown as hollow circles. The position and amino acid substitutions found in *Mm*PylRS are given for each clone.

We then performed five conjugation-editing cycles, in which we sequentially transferred the F plasmid carrying PylRS and tRNA_CUA_ genes through cells bearing each retron library. After five conjugation-editing cycles we sequenced 66 randomly selected clones from the resulting pool. This revealed that 91% of clones carried at least one edit and individual clones contained upto six edits from up to four distinct retrons, prior to any form of phenotypic selection (**Figure 4c**).

We transferred the diversified pool of F-plasmids into a selection strain harbouring a reporter plasmid. We used a sfGFP and chloramphenicol resistance dual reporter, in which an amber codon (TAG) is inserted in both genes (*cat*112TAG, *sfGFP*151TAG) to select for active synthetases, in the presence of N^ε^-benzyloxycarbonyl-L-lysine (CbzK) or N^ε^-benzyl-L-lysine (BenzK), on the basis of their fluorescence and chloramphenicol resistance. We ran these selections by plating transconjugants on plates containing chloramphenicol and the respective ncAA, and picking colonies that exhibited bright green fluorescence (**Figure 4d**). Importantly, we identified diverse solutions for each ncAA, including variants harbouring edits from up to three distinct operator libraries. We confirmed that the selected Mm PylRS variants enabled the incorporation of CbzK and BenzK into GFP by ncAA dependent GFP expression and the corresponding intact mass of purified sfGFP (**Figure 4e** and **4f, Supplementary Figure 4**).

## Discussion

We have exemplified a paradigm for genetic manipulation in which an input cargo is transferred through a series of strains (or libraries of strains) that operate on the cargo to generate an output that results from the multiplicative effect of the operations performed by each strain (or library of strains). As the cargo genetic elements are carried on an F plasmid episome that contains an oriT sequence and all the machinery for self-transmission, this approach will be compatible with essentially any specialized strain; therefore, we anticipate that our approach will be broadly compatible with diverse methods for site-specific mutagenesis and *in cell* selection^4,13,20,21^.

As oriT mediated conjugation efficiently transfers Mb-scale DNA sequences between cells^22^, we anticipate that this approach will be applicable to large genetic cargos, including the genes for biosynthetic pathways (eg: RiPPs, PKS, NRPS) where diversification is particularly challenging to realize in cells via *in vitro* library assembly methods and transformation^23,24^. The iterative use of three selectable markers enables numerous cycles of conjugation and editing with selection for transconjugants. In the examples demonstrated, the high efficiency of retron editing led to the generation of substantial libraries with library members containing up to six distinct edits and edits arising from several cycles of conjugation and editing, and these libraries were sufficient to select the phenotypes of interest. We anticipate that it will be possible to repeatedly cycle a cargo through each library of editors to achieve the full theoretical diversity of combinations in a library; therefore, the diversity generated by this approach may only be limited by the size of the cellular population used and, unlike most current approaches, is independent of the DNA transformation limit of cells. Moreover since F plasmids can be transmitted between most Gram-negative bacteria, extensions of our approach will enable the construction and selection of large libraries in other industrially- or medically-relevant microbial hosts^23,25,26^.

The core methodology we have developed here should enable the site directed mutagenesis of genes and genomic segments, to study genotype-phenotype relationships at scale and the rapid evolution of new gene function. We expect that the simple procedures developed here should be readily amenable to automation to enable the creation of very large site directed libraries on genetic elements up to the Mb-scale as a both a foundation for understanding the relationship between phenotype and genotype and for the generation of new function.

### Methods and materials

Antibiotics and arabinose were obtained from Sigma-Aldrich. Graphpad Prism version 9 was used to graph any collected data. SnapGene (version 7.2.1) and Adobe Illustrator (version 25.0.1) were used to generate illustrations of genomic operons. The pED208 was a kind gift from Prof. Gabriel Waksman and further modified by integrating the *Mm*PylRS and *Mm*^Pyl^tRNA^opt^ . The luminescent RK24 F-plasmid (pJF145) had been created previously from the RK24 Addgene #51950 plasmid^27^. The following bacterial strains were used in this work: DH10B were obtained from commercial competent cells. MDS42 *rpsL(K43R)* carrying a genomic *sacB*-*cat* were obtained from our previous published work^8^.

### Plasmid generation

Recombination and marker plasmids were obtained directly or by adapting previously published plasmids pKW20_CDFtet_pAraRedCas9_tracrRNA (tetR, p20^8^), pKW20_CDFarpa^R^_pAraRedCas9_tracrRNA (arpR, pJF164^27^), or CspRecT recombination plasmid (pFR156, pFR157 alias pFR156 *GenR*::*pheS*-*hygR* and pFR196 alias pFR156 *GenR::*Amp*R*^28^. The sequences of retron genes *Ec86* and *mrs*-*mrd* were obtained as gBlocks from IDT and cloned into a pMB1 backbone harbouring a spectinomycin resistance gene. Retron libraries were constructed by ligation of duplex synthetic oligonucleotides carrying the NNK degenerated positions with 35 bp flanking homology and overhangs complementary to a BsaI digested retron plasmid.

#### F-plasmid single step conjugation

DH10B harbouring the self-transmissible, luminescent RK24 derivate (pJF145, 50 μg/ml apramycin) or its non-transmissible derivate (pJF146, 50 μg/ml apramycin) were use as donors into a recipient MDS42 strain caring a genomic chloramphenicol resistance gene. 10 mL of each donor and recipient cultures were washed two times with PBS, conjugations were set up as a 1:1 mixture of the washed cultures with a total of 4*10^9^ cells. To initiate the conjugation the mixed cells were collected by centrifugation (5 min, 5000g), resuspended in 5–10 μL PBS and spotted on cut cellulose acetate filter membranes (Sartorius, 0.45 μm, Type 11106) placed on 2xYT agar plates. The plates were incubated at 37 °C for ∼1 h and cells were then recovered by vortexing the filter membranes in a 2 mL Eppendorf tube with 0.5 mL of SOB. The conjugation reactions were transferred to a 1.2 mL 96-well plate (AB-1127) and diluted five-fold eight times. The diluted cultures were spotted onto 2xYT agar plates selecting for the recipient cells (20 μg/mL chloramphenicol). Trans-conjugation efficiency was determined following imaging of the plates in brightfield (0.5 s) and luminescent (20 s). Images were overlayed and analysed using the Fiji software package (ImageJ, version 2.1.0/1.53c), determining the ratio of luminescent colonies over the total number of spotted recipient colonies.

#### F-plasmid iterative conjugation

DH10B harbouring the self-transmissible, luminescent RK24 derivate (pJF145, 50 μg/ml apramycin) or its non-transmissible derivate (pJF146, 50 μg/ml apramycin) were use as donors into a recipient MDS42 strain carrying a genomic chloramphenicol resistance gene. 10 mL of each donor and recipient cultures were washed two times with PBS, conjugations were set up as a 1:10 mixture of donor to recipient with overall 4*10^9^ cells.

To initiated conjugation the mixed cells were collected by centrifugation (5 min, 5000g), resuspended in 5–10 μL PBS and spotted on cut cellulose acetate filter membranes (Sartorius, 0.45 μm, Type 11106) placed on 2xYT agar plates. The plates were incubated at 37 °C for ∼1 h and cells were then recovered by vertexing the filter membranes in a 2 mL Eppendorf tube with 0.5 mL of SOB. The mixture was recovered for 30 min at 37 °C to allow for phenotypic conversion of transconjugant cells into donor cells. The next cycle of conjugation was initiated as described, this was repeated for three cycles.

In each cycle 10 μL of the conjugation reactions were transferred to 490 μL PBS in a 1.2 mL 96-well plate (AB-1127) and diluted five-fold six times. 50 μL of the diluted cultures were spotted onto 0.5xTBE agar plates (no selection) to obtain single colonies. The single colonies were imaged after 18 h at 37 °C in brightfield (0.5 s) and luminescent (20 s). Images were overlayed and analysed using the Fiji software package (ImageJ, version 2.1.0/1.53c), determining the ratio of luminescent colonies over the total number of recipient colonies in each cycle.

### Construction of a pED208 F-plasmid with integrated cat or MmPylRS and Mm^Pyl^tRNA_CUA_

As described previously in detail^29^, we utilized integration and replacement of a double-selection cassette to integrate the *MmPylRS and Mm^Pyl^tRNA_CUA_* operon on the pED208 F-plasmid. We transformed cells harbouring pED208 F-plasmid with the pKW20_CDFarp_pAraRedCas9_tracrRNA (*arpR*) plasmid expressing the lambda red alpha/beta/gamma genes, induced them with 0.5% L-arabinose, and prepared them as electrocompetent (washed twice with water or 10% glycerol). We then transformed PCR products of EM7_PheS*-HyrR double selection cassettes flanked by 50 bp homology arms replacing the copper/silver resistance operon on pED208. Following recovery, we plated cells on LB agar plates supplemented with tetracycline (5 μg/mL) and hygromycin (200 μg/mL) for *pheS*\*-*hygR*. Following confirmation of *pheS*\*-*hygR* by PCR the same process was repeated, replacing *pheS*\*-*hygR* with *MmPylRS and Mm^Pyl^tRNA_CUA_ PCR* products harbouring a kanamycin resistance gene using the same homology arms. Transformants were selected on tetracycline (5 μg/mL), kanamycin (50 μg/mL) and 2.5 mM 4-CP (4-Chlorophenylalanine), selecting for the replacement of *pheS*\*-*hygR* with *MmPylRS and Mm^Pyl^tRNA_CUA_*. The correct integration was confirmed by PCR and sequencing, this strain was used as the donor to initialize the retron editing experiments. The same procedure was used to integrate derivates of cat, utilizing a PCR product harbouring cat instead of *MmPylRS and Mm^Pyl^tRNA_CUA_*.

***Retron editing*** Cells harboring a retron plasmid, an editing helper plasmid, and the gene to be edited were grown overnight with the appropriate antibiotics added. The next day, the cultures were diluted 1:100 and grown for 1 h before induction of retron editing via the addition of 0.2% arabinose. After 3.5 to 6 h of editing, the cultures were washed and used for the next conjugation step or plated on chloramphenicol plates for determining editing efficiencies.

For conjugation-editing cycles, we immediately proceeded to a conjugation step after the editing step and selected for transconjugants overnight. The next day, we performed the next editing step followed by the next conjugation step. We iterated this process as necessary.

### aaRS library selections of F-plasmid libraries and GFP measurement

The F-integrated libraries (10 µg/mL tetracycline, 50 μg/mL kanamycine) were delivered by conjugation into DH10B cells containing the p15A-*CAT111TAG* -*sfGFP150TAG_His6_* selection plasmid (50 µg/mL Apramycine). Transconjugant cells were grown overnight and diluted 1 in 20 into selective media (20 mL) on the following day. Once cells reached exponential phase (OD_600_ >0.5) all cells were harvested and approximately 10^9^ cells were plated onto Cm selection plates (150 μg/mL Chloramphenicol, 0.2% Arabinose, 50 μg/mL Kanamycin, 10 µg/mL tetracycline, and 2 mM CbzK or BenzK, 25 x 25 cm). The plates were incubated at 37 °C for 20 h. Generally, single colonies were isolated and sequenced and the GFP expression was measured in presence and absence of the respective substrate. 50 μl of culture for each clone was diluted into 450 μl LB medium containing 0.2% L-arabinose and the appropriate antibiotics in presence and absence of 2 mM of their respective substrate. Cultures were grown in a 96 well plate with 1.2 mL wells. The wells were sealed with air permeable foil and the plate was incubated for 20 h in a shaking incubator at 37 °C, 1000 rpm. Cells were then harvested by centrifugation (3000 xg, 10 min), the growth medium discarded and the inverted plate briefly placed on paper towels (1-2 min). The cell pellets were resuspended in 150 μl

PBS, of which 100 μl resuspended cells were transferred to a clear flat-bottom 96-well plate (Nunc96). The OD_600_ and GFP fluorescence of each well was measured with the PHERAstar FS (BMG Labtech) plate reader. Mesuring OD_600_ using a 600 nm light source, and GFP fluorescence using an optical module with an excitation wavelength of 485 nm and emission wavelength of 520 nm (gain was set to 0). Plots show the average GFP fluorescence normalized by OD_600_ and the standard deviation of the triplicate measurement.

### GFP Purification and Mass Spectrometry

Cultures of F-integrated MmPylRS mutants with the p15A-*CAT111TAG* -*sfGFP150TAG_His6_* selection plasmid (50 µg/mL Apramycine) were scaled to larger volumes (0.5 mL to 10 mL LB) if necessary, and non-canonical amino acids were added at 2 mM. The cells were harvested by centrifugation and the pellet was resuspended in 1 mL of 20 mM Tris-HCl pH 8, 150 mM NaCl containing 1x BugBuster Protein Extraction Reagent, and lysed by agitating for 10 min at room temperature. The lysate was cleared by centrifugation (10 min, 15000 xg), and Ni-NTA beads (20 µL slurry) were added to the supernatant. The slurry was incubated while agitated for 1 hour (RT) and collected using a fritted spin filter (300 xg, 10 s). Following three washes with 1 mL of wash buffer (20 mM Tris-HCl pH 8, 150 mM NaCl, 40 mM imidazole) GFP was eluted in 50 µL of elution buffer (20 mM Tris-HCl pH 8, 150 mM NaCl, 200 mM imidazole). The buffer was exchanged for 20 mM Tris-HCl pH 8, 150 mM NaCl using a 10 kDa spin concentrator. High resolution mass spectra of GFP were obtained by ESI-MS using a Waters Xevo G2 MS with a modified nanoAcquity LC system, as previously reported^30,31^.Briefly, injected proteins were separated on a BEH C4 UPLC column (1.7 μm; 1.0 x 100 mm; Waters) with a flow rate of 50 μL/min using an acetonitrile gradient starting at 2% vol/vol to 80% vol/vol (0.1% vol/vol formic acid) over 20 minutes. The column outlet was directly interfaced via an electrospray ionization source with a hybrid quadrupole time-of-flight mass spectrometer (Waters). A cone voltage of 30 V was used during data acquisition in positive ion mode with a range of 300-2000 m/z. The scans were deconvoluted using the MaxEnt1 function within MassLynx software (Waters). The deconvolution program provided in the software was used to obtain the entire mass spectra. The expected, theoretical mass was calculated using ProParam (Expasy) and adapted manually for the mass difference expected for a given non-canonnical amino acid and the loss of mass during GFP maturation.

## Acknowledgments

pED208 was a gift from Prof Gabriel Waksman. This work was supported by the Medical Research Council (MRC), UK (MC_U105181009 and MC_UP_A024_1008) to JWC. MS was funded by Deutsche Forschungsgemeinschaft (DFG, German Research Foundation) SP 1981/1-1 (project no. 493404643). FBHR was supported by a UK Research and Innovation (UKRI) Marie Skłodowska-Curie Actions (MSCA) guarantee fellowship (EP/Y014154/1) and an Investigator Grant (GNT2018461) from the National Health and Medical Research Council (NHMRC) Australia.

## Author contributions

M.S., F.B.H.R., performed experiments. M.S. conceptualized iterative conjugation and editing and its applications and modifications. F.B.H.R. developed and directed efficient *in vivo* editing strategies and site-saturation library construction. J W.C. supervised the project and wrote the paper with the other authors.

## Data and materials availability

The authors agree to provide any data or materials and strains used in this study upon request, and Genbank files with DNA sequences for plasmids and genomes from this study are provided in the supplementary materials.

