## Supplementary Information for "Continuous site-directed mutagenesis and selection in *Escherichia coli*"

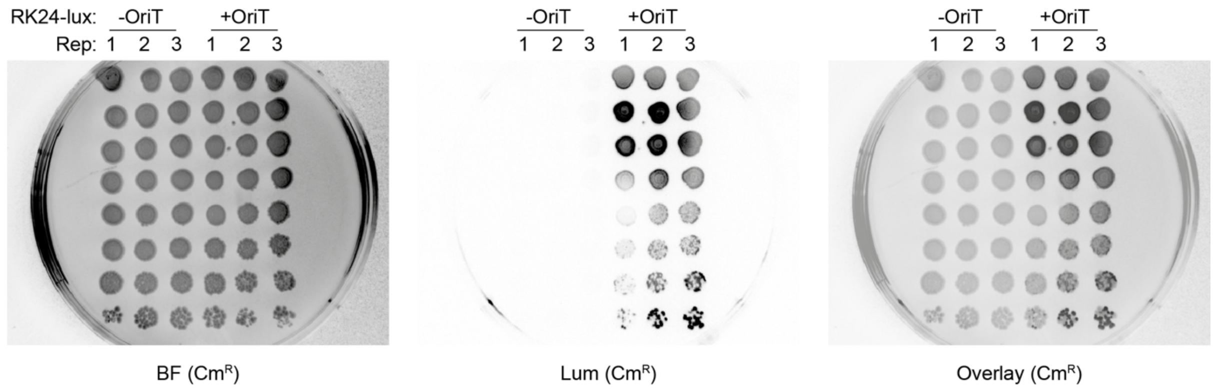


**Supplementary Figure 1**. **Single step transfer of F-integrated luxABCDE operon.** The lux operon is integrated into the RK24 F-plasmid (RK24-lux), without or with an OriT (-OriT or +OriT). Images show spotted recipient cells ($Cm^R$) following conjugation imaged as bright field (BF), luminescent (Lum) and the overlay of both (Overlay). Conjugation efficiency was determined as the ratio of luminescent single colonies over the total amount of single colonies for each replicate.

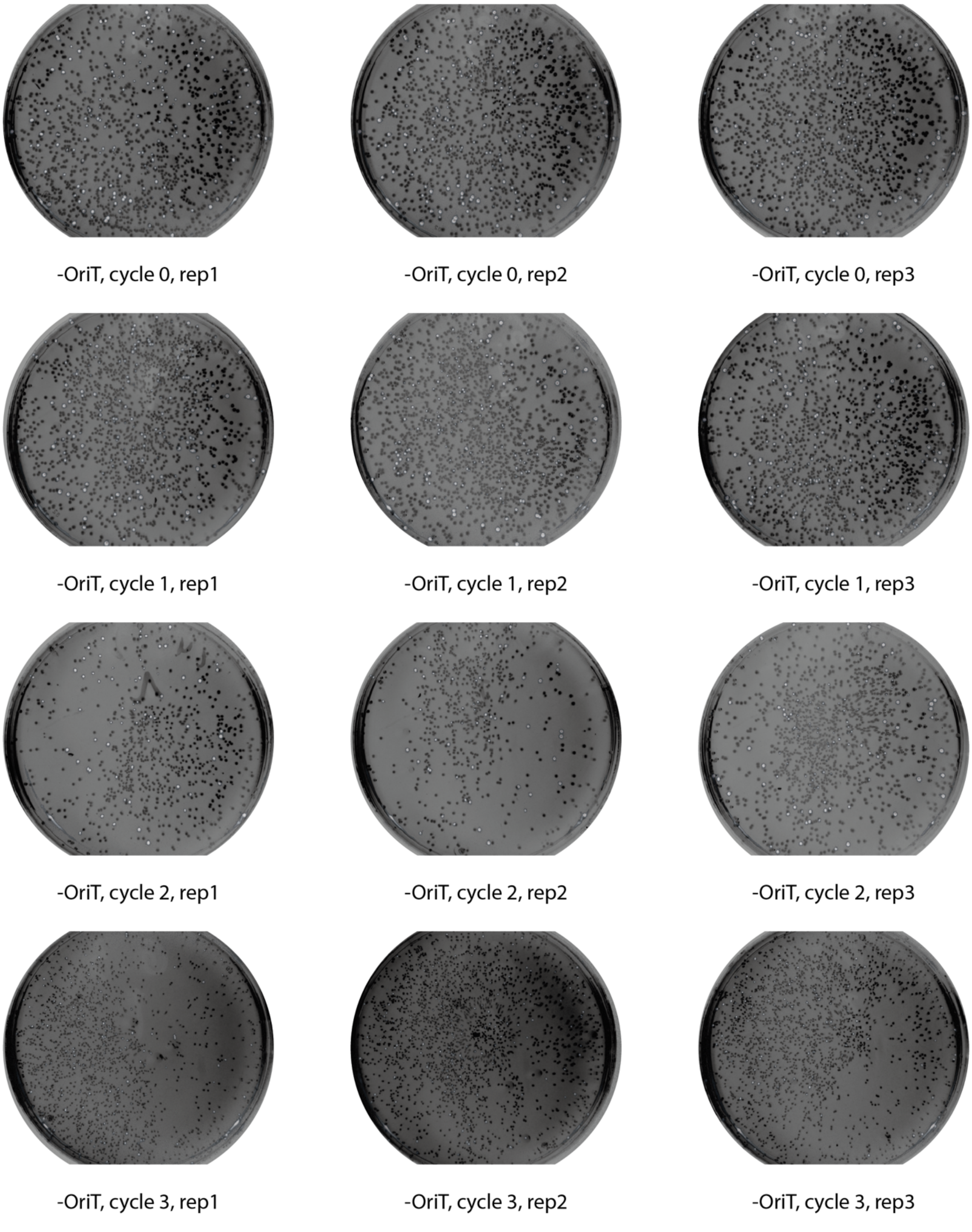


**Supplementary Figure 2**. **Colonization of a cell population by the RK24-luxABCDE F-plasmid lacking the OriT sequence (-OriT).** Images show overlayed brightfield and luminescent images of the total mixed cell population (donor and recipient, no selection) before any conjugation (cycle 0) and after 1 to 3 iterative cycles of conjugation. In the overlay, brightfield image is false coloured black and the luminescent signal is overlayed in white.

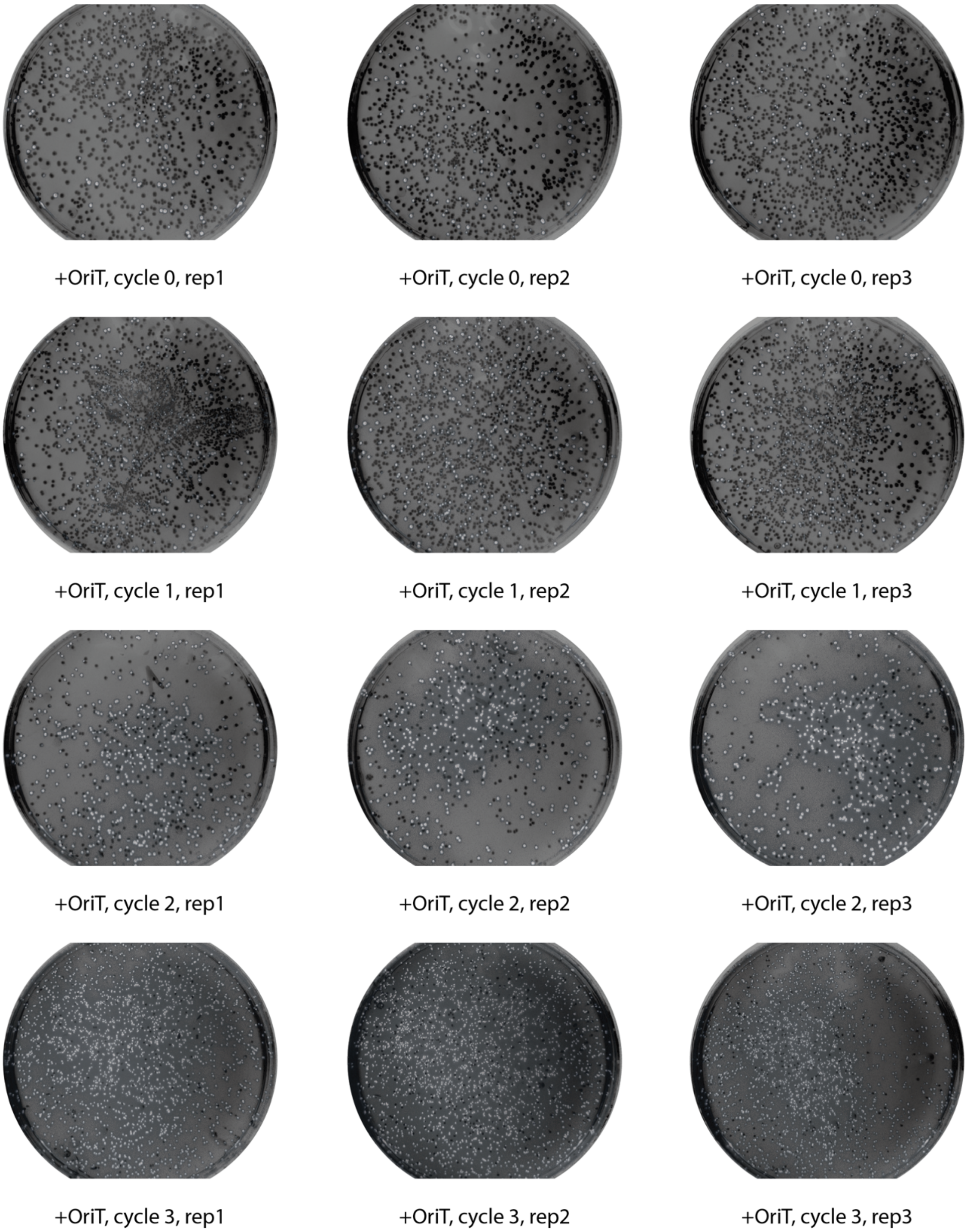


**Supplementary Figure 3**. **Colonization of a cell population by the RK24-luxABCDE F-plasmid with the native OriT sequence (+OriT).** Images show overlayed brightfield and luminescent images of the total mixed cell population (donor and recipient, no selection) before any conjugation (cycle 0) and after 1 to 3 iterative cycles of conjugation. In the overlay, brightfield image is false coloured black and the luminescent signal is overlayed in white.

Internal

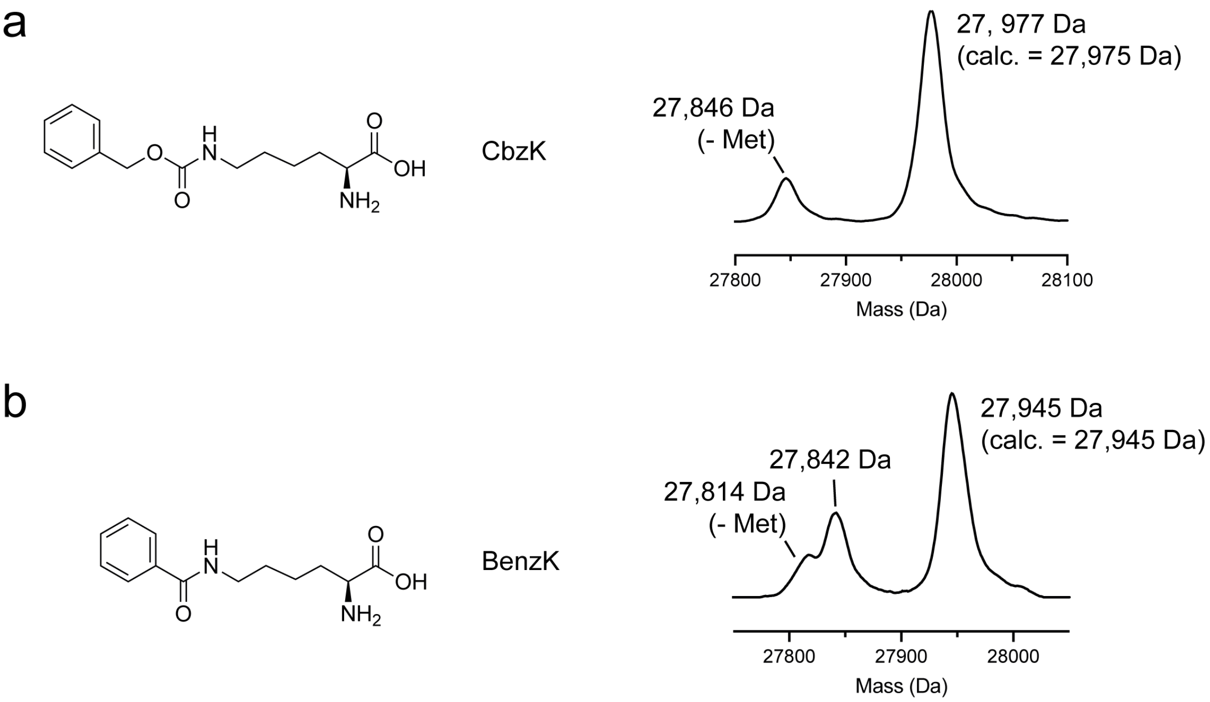


**Supplementary Figure 4**. **Analysis of GFP proteins containing a ncAA by mass spectrometry.** Shown are reconstructed mass spectra for (a) Nε-benzyloxycarbonyl-L-lysine (CbzK) or (b) Nε-benzyl-L-lysine (BenzK) in GFP. In both cases, the major species is the expected product mass.

Internal
